# Graded heterogeneity of metabotropic signaling underlies a continuum of cell-intrinsic temporal responses

**DOI:** 10.1101/2020.12.27.424473

**Authors:** Chong Guo, Vincent Huson, Evan Macosko, Wade G. Regehr

## Abstract

Many neuron types consist of populations with continuously varying molecular properties. Here we show a continuum of postsynaptic molecular properties in three types of neurons and assess the functional consequences in cerebellar unipolar brush cells (UBCs). While UBCs are thought to form discrete functional subtypes, with mossy fiber (MF) activation increasing firing in ON-UBC and suppressing firing in OFF-UBC. We find instead a continuum of response profiles that reflect the graded and inversely correlated expression of excitatory mGluR1 and inhibitory mGluR2/3 pathways. MFs coactivate mGluR2/3 and mGluR1 in many UBCs, leading to sequential inhibition-excitation because mGluR2/3-currents are faster. Additionally, we show that DAG kinase controls mGluR1 response duration, and that graded DAG kinase levels account for systematic variation of response duration over two orders of magnitude. These results demonstrate that continuous variations in metabotropic signaling can generate a stable cell-autonomous basis for temporal integration and learning over multiple time scales.

---

Throughout the brain, many cells types exhibit continuous molecular variations^1-3^. Recent studies are beginning to provide insight into some functional implications of such graded heterogeneity. Molecular diversity can reflect spatial variations^4-6^, and variation in physiological properties such as excitability and spiking properties of neurons^6,7^. Here we find that transcripts controlling expression of postsynaptic receptors and downstream signaling pathways are selectively and differentially regulated in many cell types (**Figure 1**). To evaluate the functional consequences of this graded continuum of synaptic properties, we studied cerebellar unipolar brush cells (UBCs). UBCs were recently shown to exhibit continuous molecular variations, but how this relates to physiological responses to synaptic inputs was unclear^6^. Interestingly, UBCs typically receive a single MF input and can transform brief synaptic inputs into diverse, long-lasting changes in firing^8-15^. For these reasons, UBCs are uniquely suited to dissect the relationship between transcriptomic continua and their functional implications at central synapses.

**Figure 1.**
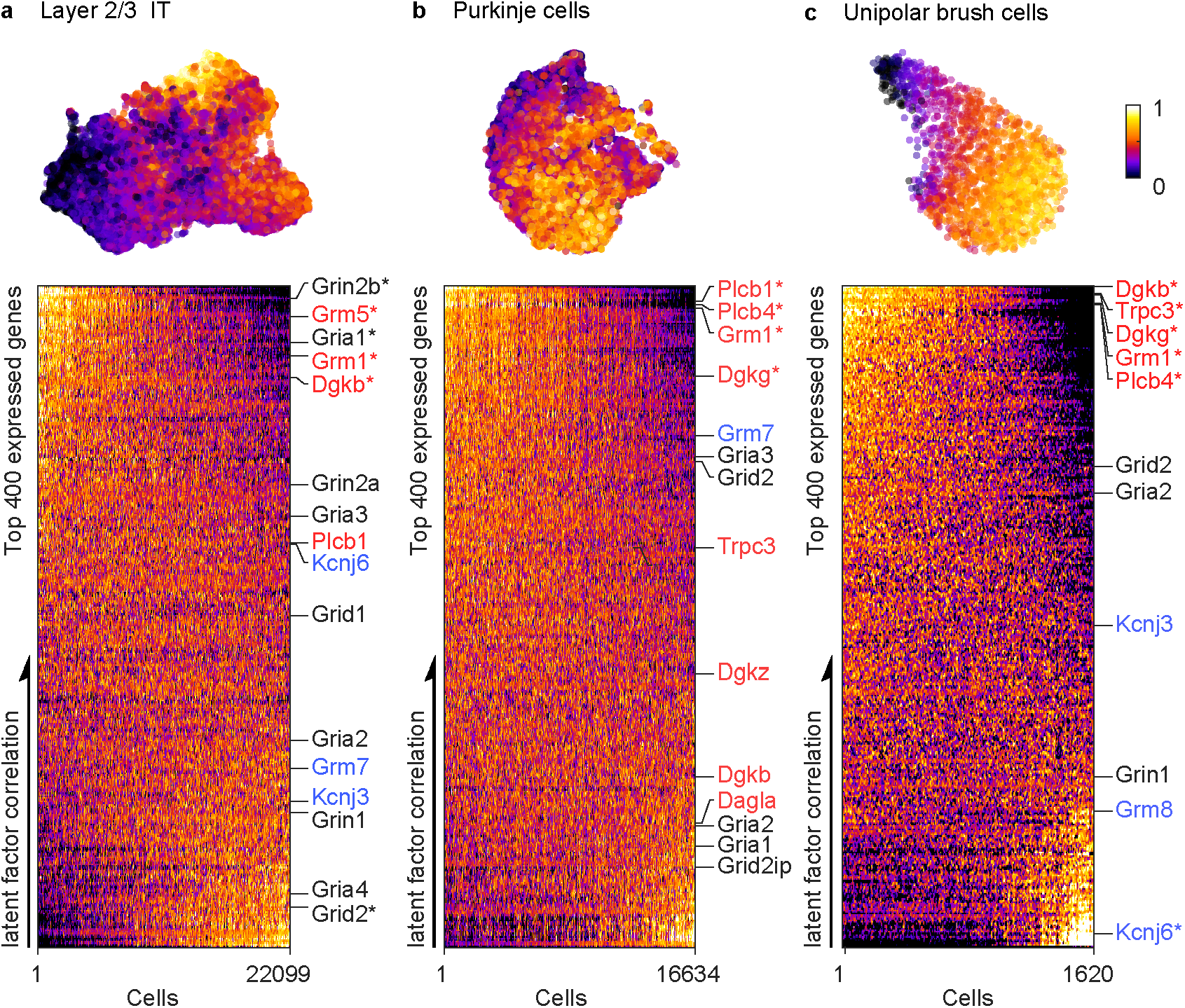
Graded continuum of molecular properties in different cell types. a. (Top) UMAP embedding showing the latent factor in cortical layer 2/3 intratelencephalic (IT) neurons. (Bottom) Heatmap of normalized gene expressions (top 400) ordered by increasing correlation with the latent factor (vertical axis). The cells are sorted in the order of decreasing factor loading (horizontal axis). All genes involved in glutamatergic synaptic transmission are shown to the right. Ionotropic receptors are shown in black, Group I metabotropic receptors and associated proteins are highlighted in red; and Group II/III metabotropic receptors and associated proteins are shown in blue. The most differentially expressed genes are denoted with an asterisk (top 1% Pearson R) b. Same analysis for cerebellar Purkinje cells. c. Same analysis for cerebellar unipolar brush cells (UBCs).

Synaptic properties of UBCs are thought to contribute to temporal learning in the cerebellum. While UBCs are present throughout the cerebellum, they are most abundant in regions controlling eye movement and in vestibular processing. To facilitate adaptative control of sensorimotor and cognitive behaviors^16-20^, the cerebellum makes learned predictions on time scales ranging from hundreds of milliseconds to tens of seconds^21^. MF inputs are transformed into high-dimensional spatial-temporal patterns of granule cell (GrC) activities at the input layer, then GrCs are pooled by the output neurons, the Purkinje cells (PCs), to produce a desired response (**Figure 2A**). The capacity of this network to learn complex input-output associations across time necessitates the existence of a diverse temporal basis set in the GrCs^22-24^. UBCs have emerged as important circuit elements within the cerebellum that can overcome the short integration time constant of GrCs. Each UBC receives a single MF input but projects onto several hundred GrCs^8^. Traditionally, UBCs are functionally divided into OFF and ON subtypes. A MF burst suppresses firing in OFF-UBCs for hundreds of milliseconds by activating inhibitory group II metabotropic glutamate receptors (mGluR2/3)^25-27^ which are coupled to GIRK2, whereas in ON-UBCs MF activation leads to a prolonged glutamate signal and long-lasting activation of AMPA receptors (AMPARs) to increase firing in ON UBCs for hundreds of milliseconds ^11^.^12^.^14^.^25^.^28^. In this way, the UBC population makes a crucial contribution to temporal representation within the input layer by preprocessing MF inputs.

**Figure 2.**
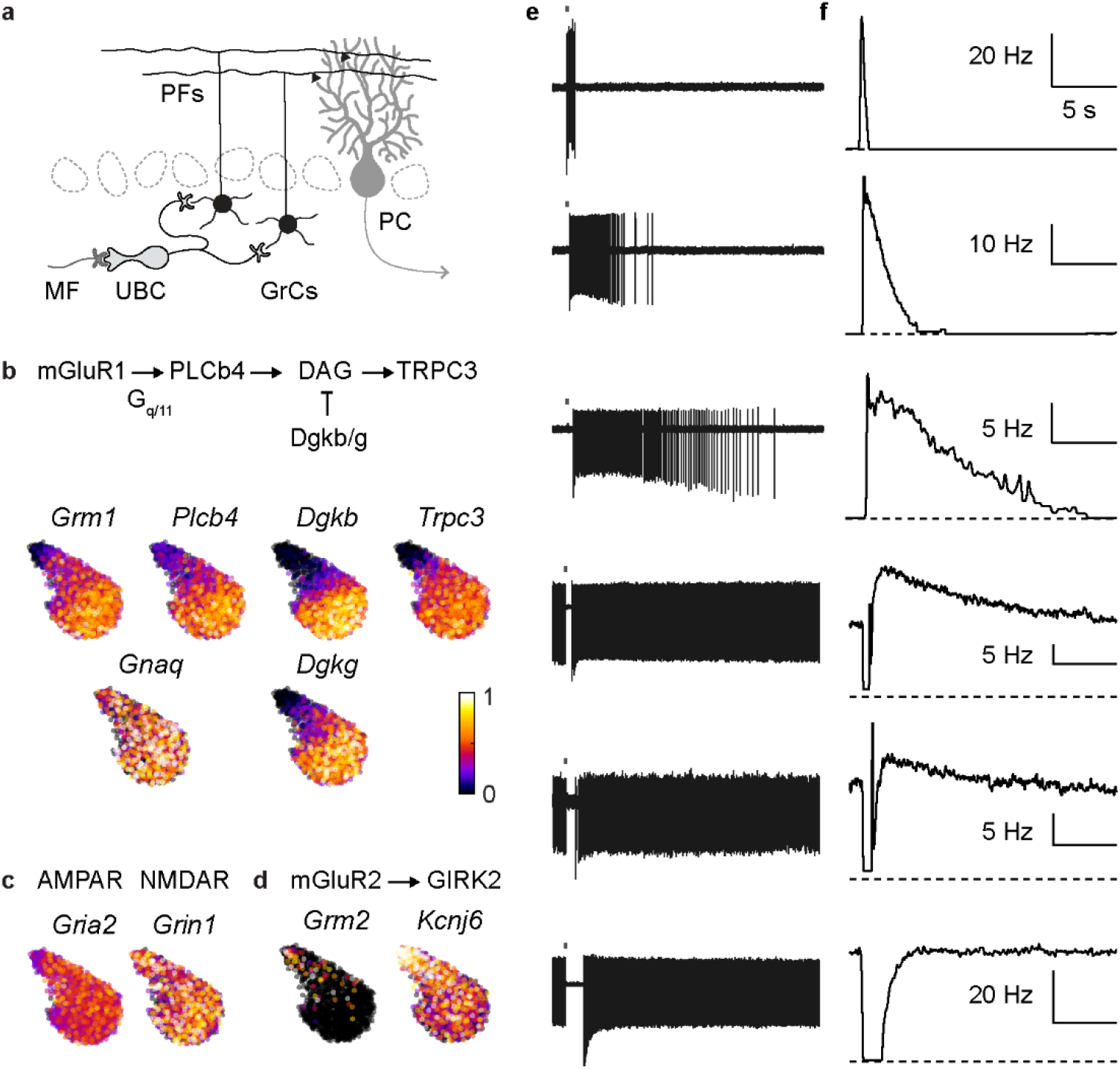
Continuous variations in molecular and functional properties of UBCs. a. Schematics of cerebellar circuit b. UMAP embedding showing normalized expression of gene involved in mGluR1 signaling cascade. c. Same as in D for AMPA and NMDA receptors. d. Same as in D for genes involved in mGluR2 signaling cascade. e. Example spiking responses in different UBCs to a burst of MF input (20 stimuli at 100Hz). f. Instantaneous firing rate for the same cells.

There are many unresolved issues regarding the generation of temporal responses in UBCs. (1) It is not known whether the diverse temporal responses of UBCs reflect factors upstream of receptor activation, such as differences in glutamate release or glutamate uptake^11,12,14,25,28^, or differences in the UBCs themselves, such as their ligand-gated receptors or their voltage-gated conductances^13^. (2) ON-UBCs are known to contain mGluR1 coupled to TRPC3 that have the potential to excite cells for longer durations than can be achieved with AMPARs, yet it is widely accepted that synaptic responses in most UBCs are dominated by AMPARs rather than mGluR1 currents^14,25,28^. (3) Molecular markers for UBC subtypes lack strict adherence to the ON and OFF dichotomy, leading some to suggest that there may be three or four subtypes of UBCs^29-31^. Notably, many UBCs contain both mGluR1 and mGluR2, but the functional implications of this are not known. (4) Single-cell RNAseq (scRNAseq) analysis suggests that UBCs have continuously varying properties, and responses evoked by pressure-applied glutamate also exhibited continuous variation over the entire population^6^, but the functional consequences of molecular diversity on responses evoked by synaptic activation are not known.

Here we reexamine the molecular and functional properties of UBCs. We observe expression gradients in components of the mGluR1 and mGluR2/3 signaling pathways that are inversely correlated, and uniform AMPAR expression. Concordantly, we find that MF activation evoked continuously varying UBC responses that ranged from brief suppression, to suppression followed by low-level long-lasting excitation, to brief intense excitation. MF excitation was mediated primarily by mGluR1 rather than by AMPARs. The amplitudes and durations of mGluR1 responses were inversely correlated and varied over two orders of magnitude, which was consistent with the coregulation of the mGluR1-TRPC3 pathway that determines amplitude, and DAG kinases that decrease response duration. We find the MFs coactivated mGluR2/3 and mGluR1 in many UBCs, with the faster mGluR2/3 component initially suppressing firing, and the slower mGluR1 component subsequently elevating firing. Together, the graded molecular variations across many components of metabotropic signaling generate a diverse continuum of cell-intrinsic synaptic responses. This provides a stable cell-autonomous mode of temporal integration and a population response which may support temporal learning over multiple timescales. Our findings reveal a surprising functional consequence of graded heterogeneity in metabotropic signaling in UBCs and suggest a possible role of such a molecular continuum in other brain areas.

## Results

Transcriptomic cell type classifications based on scRNAseq data frequently uncover discrete neuron types exhibiting continuous variations in their molecular properties^1-3^. We analyzed three such cell types to determine whether components of postsynaptic glutamatergic signaling also varied continuously in these cells. UMAP embedding of gene expression is shown for cortical layer 2/3 intratelencephalic (IT) neurons, Purkinje cells and cerebellar UBCs (**Figure 1a-c, top**). Across each of these cell types, we observed a continuous axis of molecular variation. To highlight the genes contributing to this molecular gradient, we ordered cells by their latent factor loadings (from high to low) and ordered the top 400 genes by their correlation with the latent factor (**Figure 1a-c, bottom**). We indicate the position of genes encoding glutamatergic receptors and components of downstream pathways with three different colors. Ionotropic glutamate receptors (AMPARs and NMDARs) are shown in black (*Gria1-4, Grin1, Grin2a, Grin2b, Grid1-2* and *Grid21p*). Inhibitory Group II and Group III metabotropic glutamate receptors and the inwardly rectifying potassium channels they activate in blue (*Grm2, Grm3, Grm7, Grm8, Kcnj3, Kcnj6*), and components of excitatory Group I metabotropic glutamate receptor signaling pathways in red (*Grm1, Grm5, Plcb1 Plcb4, Dgkb, Dgkg, Dgkz* and *Trpc3*). These three types of cells, layer 2/3 cortical cells (**Figure 1a**, *Grm1, Grm5*, and *Dgkb*), Purkinje cells (**Figure 1b**, *Grm1, Plcb1, Plcb4*, and *Dgkg*) and UBCs (**Figure 1c**, *Dgkb, Trpc3, Dgkg, Grm1, Plcb4*), all share the property that multiple components of excitatory group I metabotropic receptor signaling exhibit graded, continuous variations in expression levels.

The general implications of gradients in metabotropic signaling can be better appreciated by carefully analyzing the correspondence between molecular and functional properties in UBCs (**Figure 2a**). UMAP visualization of normalized gene expression for mGluR1 signaling showed similar gradients for *Grm1, Plcb4, Dgkb, Dgkg*, and *Trpc3*, but *Gnaq* expression was uniform (**Figure 2b**). NMDARs and AMPARs are both present in UBCs. Yet while AMPA receptors were thought to contribute to diverse excitatory responses of UBCs^11,12,14,25,28^, genes encoding AMPARs (*Gria2*) and NMDARs (*Grin1*) were not differentially expressed (**Figure 2c**). Genes encoding mGluR2/3 (*Grm2/3*) and the downstream effector GIRK2 (*Kcnj6*) also exhibited graded expression across the population (**Figure 2d**). The transcriptomic level co-regulation of mGluR1 and mGluR2/3 signaling pathways could potentially alter the relative strength and kinetics of synaptically-evoked metabotropic currents across the UBC population.

We assessed the functional consequences of these molecular properties by examining MF-evoked responses in a large population of UBCs. Cell-attached recordings of UBCs were performed in acute slices of adult mouse cerebellum in the presence of inhibitory synaptic blockers (see Methods). Brief electrical stimulations (20×100Hz) of a MF input elicited a unique spiking response in each cell. Across cells, there was a tremendous diversity in the duration, amplitude, and sign of MF-evoked responses (**Figure 2e**). In the extremes, MF activation either evoked a large transient increase in firing (**Figure 2e**, top cell), or transiently prevented firing (**Figure 2e**, bottom cell), in line with binary ON and OFF functional classification. However, most UBCs exhibited graded responses that did not readily conform to these two categories. Excitatory responses had highly variable durations in firing that persisted from hundreds of milliseconds to tens of seconds–a difference of two orders of magnitude. This is illustrated by the spiking evoked in four different cells (**Figure 2e**, top four cells), for which the half-decay times of the instantaneous firing frequencies were 410 ms, 1.8 s, 6.8 s and 8.8 s respectively (**Figure 2f**, top four cells). There was considerable variability in the spontaneous activity of UBCs, but for cells that had fired spontaneously at high frequencies, MF stimulation tended to transiently suppress firing (**Figure 2f**, bottom three cells). For many of these cells (**Figure 2e**, cells four and five), MF stimulation evoked a biphasic response profile where a pause in firing was followed by a small but long-lasting increase in firing. Importantly, the temporal profiles of spiking responses were cell-specific and did not depend on either the intensity (**Extended Data Figure 1**) or the number of stimuli (**Extended Data Figure 2a, b**).

The large systematic variations in the amplitudes and durations of excitation are surprising. Previous studies suggested that MF-evoked excitatory stimulation generated diverse responses which were primarily the result of prolonged glutamate elevation and AMPAR activation, but these responses typically lasted for hundreds of milliseconds^14,25,28^. While others have observed slow mGluR1 responses in some UBCs, these cells were excluded from further characterization ^28^. Yet the scRNAseq data suggest that the functional gradient is potentially related to cell-intrinsic variations in mGluR1 signaling instead of the uniform expression of AMPARs or NMDARs.

To examine the relative contributions of ionotropic and metabotropic glutamate receptors to synaptically-evoked responses, we determined the effects of receptor antagonists on MF-evoked spiking. The co-application of AMPAR and NMDAR antagonists had negligible effects on MF-evoked firing in either fast or slow spiking UBCs (**Figure 3a-b**). In contrast, an mGluR1 antagonist strongly attenuated MF-evoked firing in every cell regardless of decay kinetics (**Figure 3c-d**). Thus, mGluR1 activation underlies most synaptic excitation of UBCs, and their responses kinetics are diverse, lasting from hundreds of milliseconds to tens of seconds.

**Figure 3.**
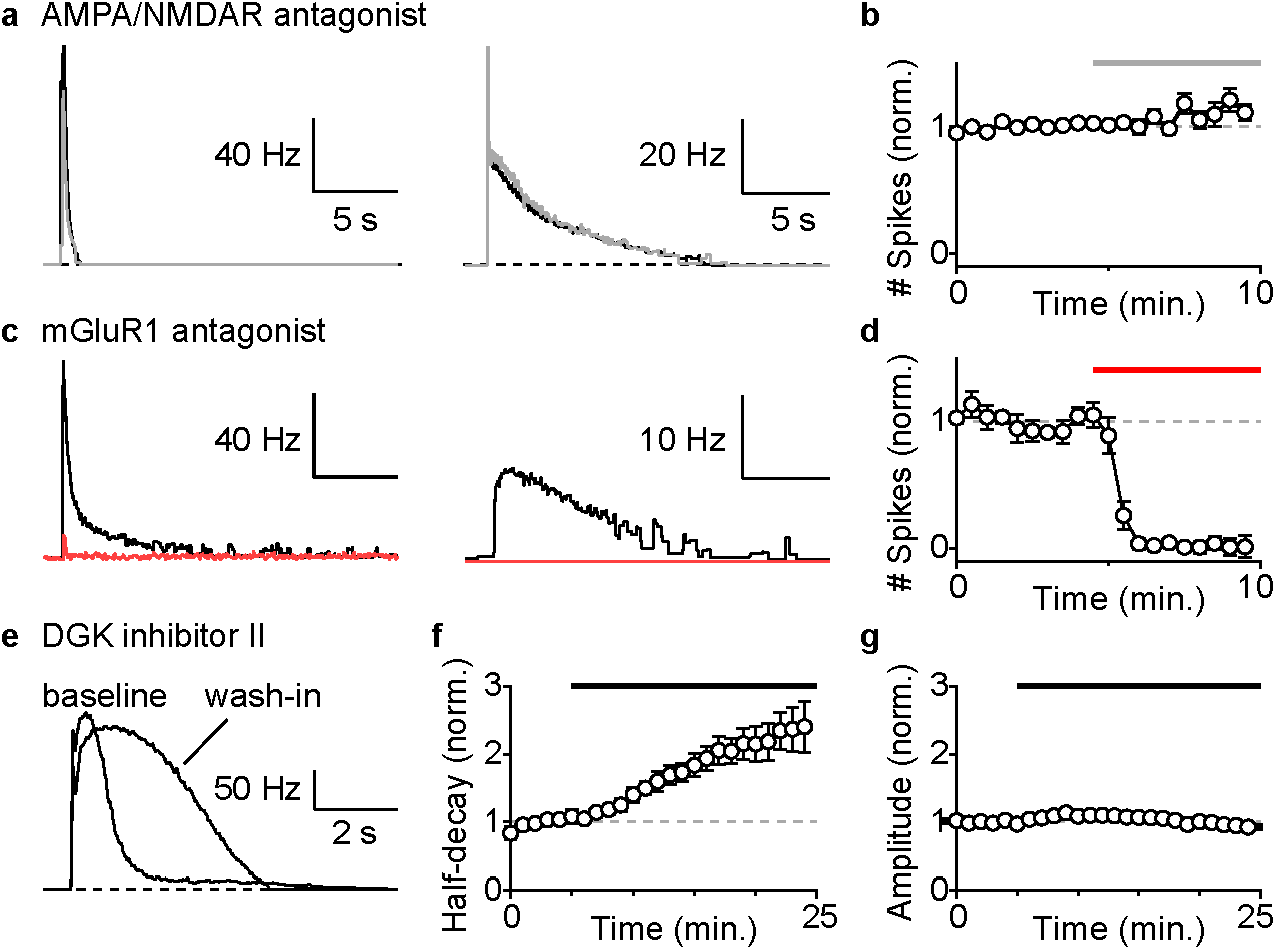
mGluR1 and DGK control excitatory mossy fiber responses in UBCs. a. Examples of instantaneous firing rate from on-cell recordings before (black) and after (gray) the application of AMPA/NMDA receptor antagonist (gray bar) in a fast (left) and a slow (right) UBC. Each trace is an average of 8 trials. b. Summary of evoked spiking (20×100 Hz) and the effect of AMPA/NMDA receptor antagonist (normalized to baseline, mean±sem, n=5). c. Examples of instantaneous firing rate from on-cell recordings before (black) and after (red) the application of an mGluR1 antagonist (red bar) in a fast (left) and a slow (right) UBC. Each trace is an average of 8 trials. d. Summary of evoked spiking (20×100 Hz) and the effect of an mGluR1 antagonist (normalized to baseline, mean±sem, n=5). e. Example of instantaneous firing rate before and after 20 minutes of DGK inhibitor II wash-in. f. Summary of half-decay time of instantaneous firing rate response with DGK inhibitor II (normalized to baseline, mean±sem, n=6) g. Summary of peak amplitude of instantaneous firing rate response with DGK inhibitor II (normalized to baseline, mean±sem, n=6)

DGKβ/γ are thought to terminate DAG signaling, but their involvement in mGluR1-dependent synaptic responses in the nervous system is not clearly understood. The concomitant rise of DGKβ/γ expression in UBCs with higher mGluR1 levels (**Figure 2b**), suggests a possible role of DGKβ/γ in regulating the decay kinetics of mGluR1-dependent spiking responses. We tested the effects of DGK inhibitor II on mGluR1-dependent responses with AMPAR antagonist in baseline condition. Bath application of DGK inhibitor II increased the half-decay time of the spiking response in UBCs (**Figure 3 e, f**, n = 6) without changing peak response amplitude (**Figure 3g**). No change in spiking response kinetics was observed in control wash-in experiments (**Extended Data Figure 3**). This suggests that the cell-intrinsic differences in DGK expression influences the variable duration of sustained spiking, with higher levels of DGK corresponding to shorter mGluR1-dependent synaptic currents and faster decays of UBC spiking responses.

The time courses of UBC firing could be controlled by the duration of mGluR1-mediated synaptic currents, but active amplification could prolong the influence of otherwise transient synaptic currents^32^. We tested for a contribution from active amplification in cells for which MF activation evoked long-lasting increases in firing. In current-clamp, transient depolarization of UBCs with current injections elevated firing without evoking persistent activity (**Figure 4a**). We also found that for MF-evoked responses in the same cell, suppressing the initial spikes with a hyperpolarizing current did not prevent the subsequent persistent firing (**Figure 4b**). Finally, we tested whether transient hyperpolarization triggered subsequent long-lasting increases in firing, but we found that the evoked rebound spiking was much smaller and shorter-lived than MF-evoked spiking (**Figure 4c**). These experiments suggest that sustained MF-evoked spiking responses in UBCs do not require active amplification.

**Figure 4.**
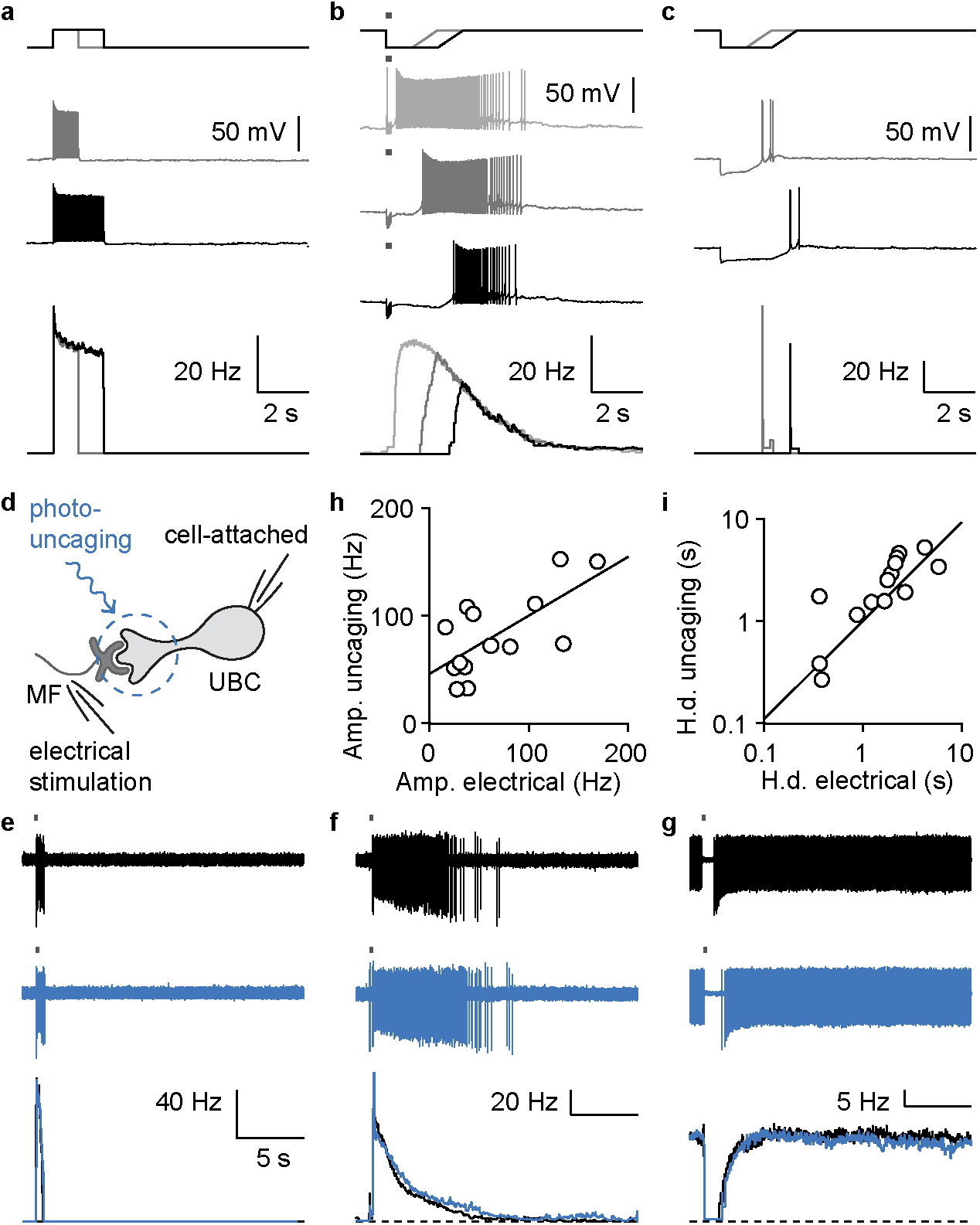
Diversity in firing following mossy fiber activation does not result from voltagedependent active amplification but reflects intrinsic synaptic differences in UBCs. a. Sample spiking responses in current clamp to 1s (grey) or 2s (black) current injections and their corresponding firing rates (bottom). b. Sample spiking responses in current clamp to MF stimulation alone (light grey) or in conjunction with a 1s (dark grey) or 2s (black) hyperpolarizing step and their corresponding firing rates (bottom). c. Sample spiking responses in current clamp to 1s (light grey) and 2s (dark grey) hyperpolarizing steps and their corresponding firing rates (bottom) d. Schematics of the electrical stimulation and optical glutamate uncaging experiment e. Example spiking response to electrical (black, top) or optical stimulation (blue, middle) and their instantaneous firing rate (bottom) for a fast UBC f. Same as in B but for a slow UBC g. Same as in B but for a UBC that was suppressed by MF stimulation h. Correlation between the amplitudes of the glutamate-uncaging and electrical spiking responses, linear fit (R_adj_^2^= 0.33, slope = 0.42, intercept = 52, n = 14). i. Correlation between the half-decay times of glutamate-uncaging and electrical spiking responses, linear fit on log_10_ transformed data (black, R_adj_^2^= 0.51, slope = 0.92, n = 14).

These diverse synaptic responses could reflect differences in either cell-intrinsic properties or extrinsic factors such as the presynaptic release properties or feedforward UBC connections. To quantify cell-intrinsic influence, we measured the kinetics of the spiking response to both electrical stimulation and optically uncaged glutamate (see Methods). We found optically uncaged glutamate recapitulated electrically evoked responses remarkably well (**Figure 4d-i**), reproducing the entire spectrum of UBC responses. In a cell with a brief MF response, uncaged glutamate mimicked the rapid kinetics of the spiking response (**Figure 4e**). Likewise, in a cell where firing persisted for many seconds after the electrical stimulation, uncaged glutamate resulted in similarly prolonged firing (**Figure 4f**). Lastly, MF stimulation and uncaged glutamate both transiently suppressed firing (**Figure 4g**). The half decay times (**Figure 4h**), and the amplitudes of spiking (**Figure 4i**) evoked by uncaging and MF stimulation were linearly correlated. Importantly, altering the glutamate release by varying the duration of the light-pulse had minimal effect on the decay kinetics (**Extended Data Figure 2d-f**). This suggests that the heterogeneity in the responses of different UBCs primarily reflected differences in cell-intrinsic properties of metabotropic signaling.

To test whether the time courses of the instantaneous firing rate closely reflected the underlying synaptic currents, we recorded MF-evoked responses in each UBC in both cell-attached and whole-cell voltage-clamp configurations. UBCs that were briefly excited had rapidly decaying synaptic currents (**Figure 5a**), and as the duration of spiking increased the synaptic currents became slower (**Figure 5b, c**). MF stimulation briefly suppressed firing in many UBCs, even in those that had a subsequent increase in firing (**Figure 5d**). For such cells, a brief outward current preceded a long-lasting but smaller inward current. In cells without spontaneous activity, the outward current produced a noticeable delay in spike initiation (**Figure 5c**). In spontaneously active UBCs, MF stimulation evoked an approximately one-second pause in firing, and in whole-cell recordings, the same MF stimulation evoked an outward synaptic current lasting about a second (**Figure 5e**). The half-decay times of the spiking increases and the synaptic currents were correlated (**Figure 5f**, R^2^ = 0.91, slope = 0.88, intercept = -0.22), as were the peaks of the firing rates and magnitudes of the inward currents (**Figure 5g**, R^2^ = 0.69, slope = 0.71, intercept = -0.65). We found that the current typically outlasted spiking by a factor of 2.0±0.6 (n=17). This was consistent with a supra-linear f-I curve which temporally sharpened the spiking response. Similarly, the amplitudes of the outward current correlated well with the duration of spike suppression (**Figure 5h**). These observations suggest that the amplitudes and time courses of synaptic currents are the main determinants of the magnitude and kinetics of firing rate increases.

**Figure 5.**
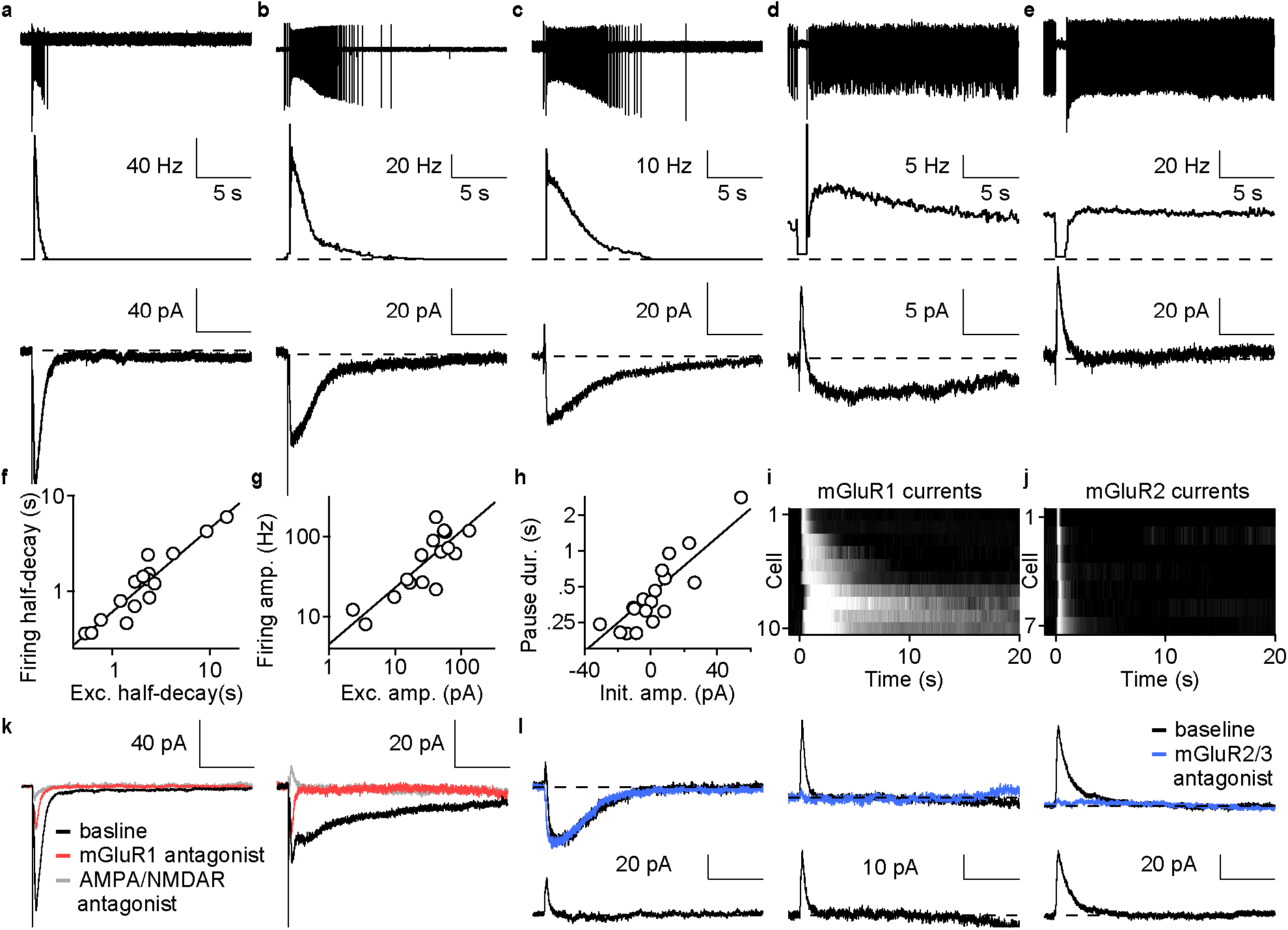
mGluR1 and mGluR2-mediated synaptic currents determine the spiking kinetics. a. Sample cell-attached recording (top) instantaneous firing rate (middle) and synaptic current measured with whole-cell voltage clamp (bottom) in a cell with fast response. b. Same as in A but for a cell with intermediate speed response c. Same as in A but for a cell with clear biphasic synaptic current (bottom) d. Same as in A but for a cell with slow biphasic response e. Same as in A but for a cell with only a pause in firing f. Half-decay times of firing rates vs. half-decay times of currents and linear fit on a log-log plot (R_adj_^2^= 0.91, slope = 0.88, intercept = -0.22, n = 17) g. Peak firing rate vs. peak current amplitude and linear fit on a log-log plot (R_adj_^2^= 0.69, slope = 0.71, intercept = -0.65, n = 17) h. Pause duration vs. amplitude of the current at stimulation offset, and linear fit with log_10_ response variable (black, R_adj_^2^ = 0.44, slope = 0.01, intercept = -0.42, n = 20) i. Heatmap of peak normalized mGluR1-mediated current (n=10) j. Heatmap of peak normalized mGluR2/3-mediated current (n=7) k. Average synaptic response before (black), after (red) mGluR1 antagonist, and after AMPA/NMDA receptor antagonist wash-ins (grey). Each trace is an average of 8 trials. l. Average synaptic response before (top, black) and after (top, blue) mGluR2/3 antagonist washin, and their difference (bottom, black). Each trace is an average of 8 trials.

To examine the contribution of mGluR1 to synaptic currents in the MF-UBC synapse, we carried out whole-cell voltage-clamp recordings. We found that mGluR1 responses were prone to washout. We therefore always identified MF inputs before break-in and used a freshly prepared K-Methanesulfonate based regenerating solution to provide the stability required for pharmacology (**Extended Data Figure 4**, see Methods). Sequential wash-ins of an mGluR1 antagonist followed by AMPAR/NMDAR antagonists revealed that most synaptic current was mediated by mGluR1 (**Figure 5k**). mGluR1 antagonists eliminated 91±5% (n = 6) of the total synaptic charge while subsequent wash-in of AMPA/NMDA receptor antagonists eliminated only a small, rapid component of the synaptic current (**Figure 5k, Extended Data Figure 5a).** The large cell-to-cell variations in the time course of synaptic currents before drug wash-in were eliminated after blocking mGluR1 (**Extended Data Figure 5b**). These observations indicate that MFs evoke increases in UBC spiking primarily by activating mGluR1s, and that diversity in the kinetics of mGluR1 currents is the primary determinant of the diverse spiking time courses.

To understand how mGluR2/3 currents relate to mGluR1 currents in UBCs, we used an antagonist to isolate the mGluR2/3 component. mGluR2/3 currents always activated during the onset of mGluR1 current in biphasic responding UBCs (**Figure 5l**, left) and mediated all the outward currents that suppressed spiking (**Figure 5l**, middle and left). The amplitude and time course of mGluR2/3-mediated current was variable in different cells (**Figure 5l**). The mGluR2/3 component was more rapid than the mGluR1 component and terminated before peak mGluR1 response (**Figure 5i, j**). These experiments suggest that the outward mGluR2/3 current makes important contributions to the initial temporal dynamics by suppressing spiking for variable durations in different UBCs.

MF stimulation evoked remarkably complex and diverse responses in UBCs (n=84). Given that molecularly, UBCs were well-described along a single axis of variation, could there be a simple organizing principle for the functional properties of UBCs at a population level? We used the half-decay time of the firing rate elevation to sort most UBCs, and we used pause duration to sort the remaining UBCs for which firing was primarily suppressed (**Figure 6a, b**). For visualization, we normalized the first group of cells by the peak firing rate increases (n=70), and the second group by baseline firing rate (n=14). The sorted heatmap plotted on a linear time axis (**Figure 6a**) and on a logarithmic time axis (**Figure 6b**), along with summary plots (**Figure 6c-h**) revealed the continuous nature of the population response and several intriguing distributional properties. For cells that responded with elevated firing, their peak times and half-widths varied continuously over two orders of magnitude (**Figure 6c** peak time, R_adj_^2^ = 0.96, slope = 0.027, intercept = -1.50, **6d** half-width, R_adj_^2^ = 0.99, slope = 0.026, intercept = -0.86). The half-width to peak-time ratio was constant across all cells (**Figure 6e**, HW-PT ratio = 4.2±0.2, n = 70) which resulted in constant width of temporal tuning curves across cells on a logarithmic time scale (**Figure 6f**, H, R_adj_^2^ = -0.015, slope = 0.0, intercept = 14.5). The distributions of the logarithm of half-widths and peak-times were indistinguishable from a uniform distribution (KS-tests, p = 0.793 and 0.9473), which were reflected in the uniform progression of peak times on a logarithmic heatmap (**Figure 6b**). Pause durations from all cells, including those in which suppression dominated, followed a linear trend on the logarithmic time scale (**Figure 6g**, R_adj_^2^ = 0.78, slope = 0.027, intercept = -2.22). Lastly, the maximum firing rate amplitude saturated above 200 Hz and then decreased nearly linearly on the log plot (**Figure 6h**, R_adj_^2^ = 0.95, see Methods). Small increases in firing were also observed in the predominantly suppressed UBCs and they followed the same trend (**Figure 6h**, grey markers). The continuity of pause and amplitude distributions strongly suggests that UBCs comprise a single population with a continuum of functional properties. A remarkable property of UBC responses is the tight tradeoff between amplitude and half-width of decay, potentially reflecting the correlated expressions of mGluR1 and DGK (**Figure 2b**). This is readily appreciated by overlaying the un-normalized responses from three representative UBCs (**Figure 6i**). Additionally, we found that every UBC response was well-approximated by shifting a single base Log-Gaussian response curve (**Figure 6i, j**, black traces). This means that on a log-log plot, UBC responses appeared as quadratic curves with shifted peak amplitudes and peak times, but constant widths (**Figure 6j**). The amplitude and duration trade-off gave rise to an approximate conservation of the average spiking response in each UBC. This was reflected in the shifted power-law decay of the population response regardless of the number of stimuli (**Figure 6k, l**, R_adj_^2^ = 0.99, see Methods). We found that UBCs were able to act as single-neuron signal integrators. While repeated MF stimulations resulted in a succession of bursts in a fast UBC (**Figure 6m**), UBCs with slower kinetics summated the input over time (**Figure 6n, o**). Responses to sinusoidally modulated MF inputs produced a range of phase delays from cell-to-cell, which could serve an important role in adaptive vestibular behaviors (**Extended Data Figure 6**). Taken together, the continuum of response properties allows the UBC population to function both as a distributed array of neural integrators with high memory capacity^33^, and as an efficient basis set for learning at multiple time-scales (**Figure 7c,d**).

**Figure 6.**
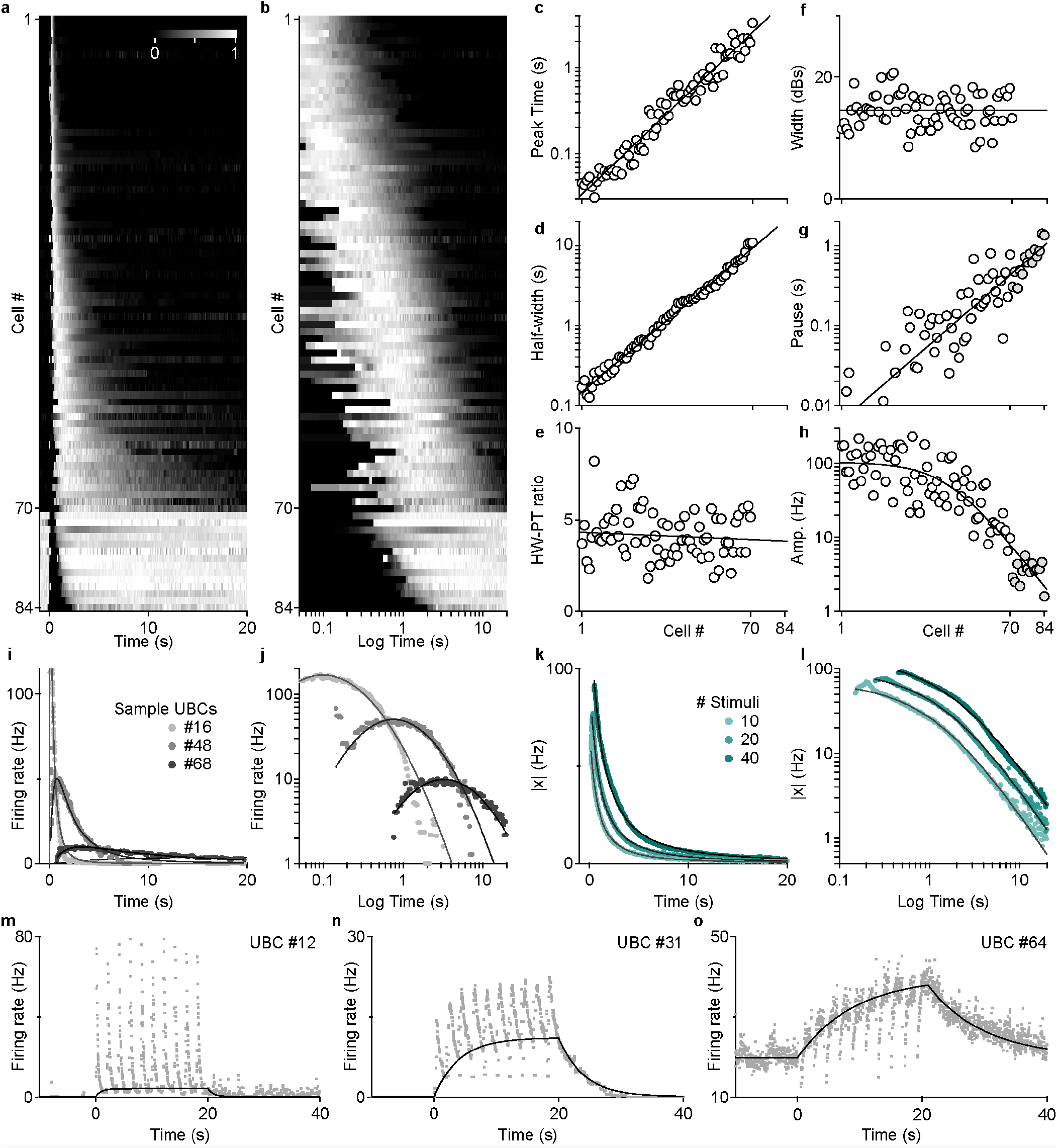
UBC population generates a continuum of multi-scale temporal representations. a. All normalized responses displayed as a sorted heat map in linear time (n = 84). b. The same sorted heat map displayed in logarithmic time (t=0 at stimulus offset). c. Peak-time vs. cell #, linear fit on the log_10_ data (black) d. Half-width of response vs. cell #, linear fit on the log_10_ data (black) e. Half-width to peak-time ratio vs. cell #, linear fit (black) f. Width of Gaussian fits of responses on a logarithmic time scale vs. cell #, linear fit (black) g. Pause time vs. cell #, linear fit on the log_10_ transformed data (black) h. Amplitude of response vs. cell #, rectified linear fit on the log_10_ data (black) i. Sample instantaneous firing rate in three UBCs (grey dots) and log-Gaussian fits (black) j. Same as in I but in a log-log plot k. Amplitude of population response over time for different numbers of stimuli (turquoise dots, 10x, 20x, and 40×100 Hz) and shifted power-law fits (black lines) l. Same as in K in a log-log plot to better illustrate the power-law decay in the tail, for 10, 20, and 40 stimuli. The critical exponent α = -1.47 with 95% CI of (-1.51, -1.44). m. Representative response of a fast UBC to a 10×0.5Hz sequence of 10×100Hz MF bursts n. Response of an intermediate UBC to the same sequence of MF bursts o. Response of a slow UBC to the same sequence of MF bursts.

**Figure 7.**
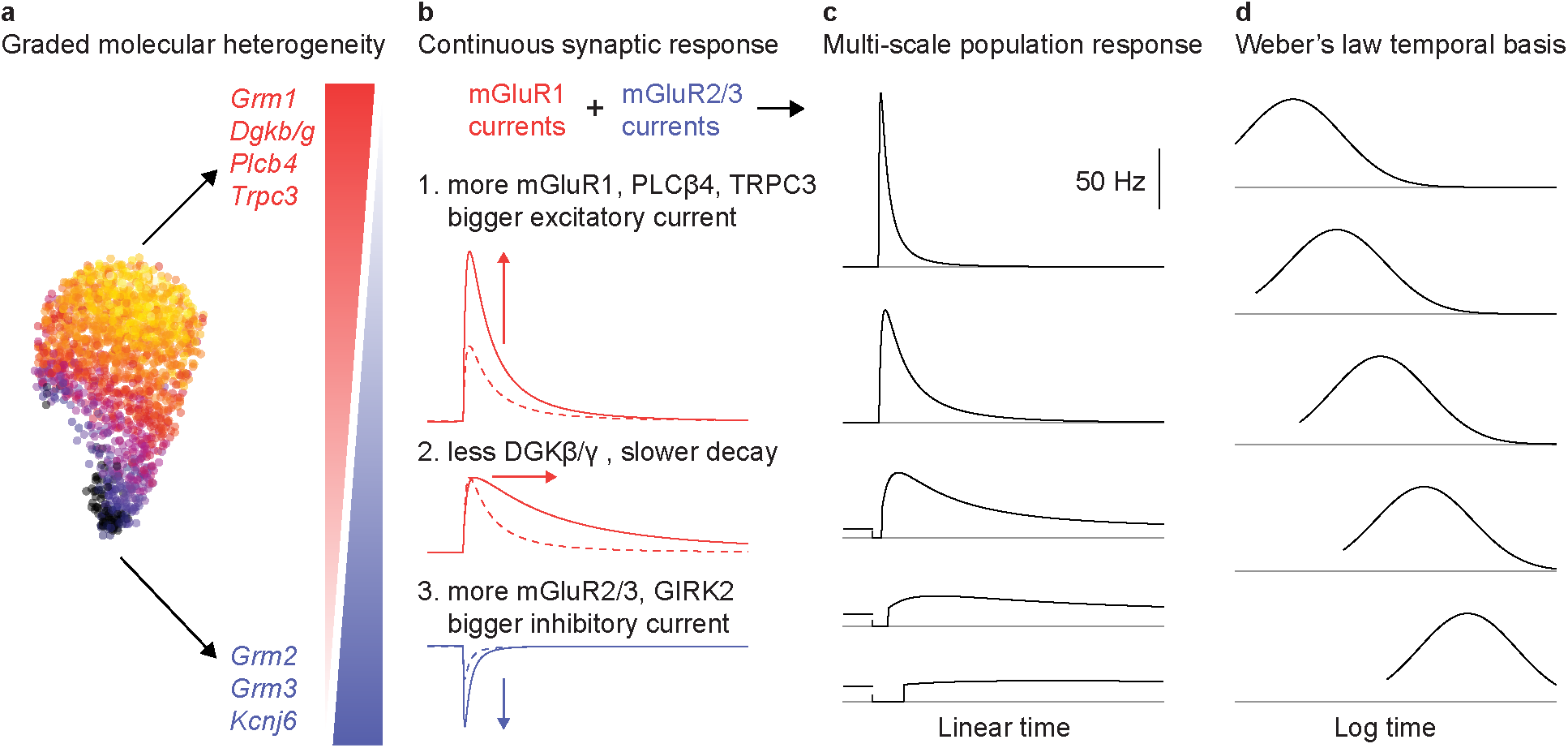
Schematic showing the roles of mGluR2, mGluR1 and DGK in controlling responses evoked by MF activation. a. UMAP with latent factor loading (left) and major genes involved in synaptic transmission that are differentially regulated along the gradient of the latent factor (right). b. Schematics summarizing the role of the mGluR1 pathway, including DGKβ/γ (red) and the mGluR2/3 pathway (blue) in continuous changes in synaptic responses. c. Schematics of temporal responses from a small group of UBCs showing the multiscale nature of temporal representation at a population level. d. Same five cells normalized and replotted on a logarithmic time axis clearly demonstrate the Weber’s law behavior of the UBC population temporal basis and its ability to support learning over multiple time-scales.

## Discussion

Our main finding is that the population of UBCs provides a continuum of cell-intrinsic temporal transformations over an exceptionally wide range of time scales (**Figure 6**), and that these transformations arise from the continuous variations in the amplitudes and time courses of metabotropic signaling, that in turn control the extent and the duration of the suppression and excitation of firing (**Figures 3-5, 7b**). These features are consistent with the observed graded molecular heterogeneity in metabotropic signaling pathways (**Figures 2, 7a**), and it is likely that similar gradients in mGluR1/5 expression have similar consequences in other cell types (**Figure 1**).

### New insights into excitatory responses in UBCs

Our findings are a significant departure from previous population-level descriptions and mechanistic accounts of temporal processing in UBCs. We have shown that the diverse temporal responses of UBCs reflect cell-intrinsic synaptic differences rather than differences in glutamate release or glutamate uptake. We have also shown that MF excitation of UBCs is mediated primarily by activation of mGluR1, in contrast to previous studies that emphasized the importance of AMPAR signaling and concluded that MF activation of mGluR1 was less prominent^14,25,28^. There are a number of possible explanations for these differences. First, some studies measured UBC responses in weakly electric fish^15^ and the auditory system^25^, where it is possible that AMPAR signaling allows more rapid excitation of UBCs tailored to the temporal demands of those regions. For example, in weakly electric fish excitatory UBC responses persist for about 10 ms to 100 ms, which is a time scale suited to cancelling very brief signals associated with the electric organ discharge^15^. Another contributing factor may be that most previous studies used whole cell recordings to characterize UBC responses^11,12,14,25,28^, and as we have shown mGluR1 responses can wash out under such conditions. Finally, one study selected cells that had simple classic ON and OFF UBC responses, and elected to not include cells with prolonged responses that did not conform to these categories^28^.

The systematic variation over two orders of magnitude in the amplitudes and time courses of MF-evoked mGluR1 responses within a single cell type was unprecedented. Previous studies suggested that glutamate diffusion determines the time course of excitatory responses by influencing the magnitude and duration of AMPAR activation^11,12,14^. Our caged glutamate experiments and the close correspondence between the amplitudes and time courses of MF-evoked and caged-glutamate-evoked excitation indicate that the properties of metabotropic signaling in UBCs are the primary determinants of the time courses of excitatory responses (**Figure 4d-f**). We also found that although UBCs have active conductances that can result in rebound firing, MF-evoked increases in firing are dominated by the mGluR1-mediated currents (**Figure 4a-c**).

Another notable feature of excitatory responses was that short-lived responses were at higher frequencies, such that MFs evoked a similar total number of spikes in UBCs regardless of the amplitude or duration of the responses. This is a natural consequence of the correlated expression of elements of the mGluR1 pathway. mGluR1 activation engages G proteins which in turn activate PLCβ4, elevating DAG levels that ultimately activate TRPC3. The responses are larger when the expression levels of mGluR1, PLCβ4, and TRPC3 are higher. In addition, the larger responses are associated with cells that have high DGK levels, rapid degradation of DAG and short-lived TRPC3 activation (**Figure 7b**). There is continuous variation in the amplitudes and time course of responses in UBCs that conform to the classic ON-UBC category, but mGluR1 responses become particularly small and long lasting in UBCs that also have a prominent mGluR2/3 component.

### mGluR2/3-mediated pauses in firing

mGluR2/3 strongly influenced the firing of a surprisingly large fraction of UBCs. MFs simply suppress firing in 17 % (14 out of 84) of UBCs, and these cells would traditionally be categorized as OFF UBCs. Even in this population of cells, the duration of suppression is graded and exhibits continuous variation. Previous studies had observed UBCs with mGluR2/3 and mGluR1 components, but here we see the functional importance of the presence of both types of receptors in the same cells. We see an inhibitory mGluR2/3 component in an additional 50% (42 out of 84) of cells that also have long lasting excitatory responses. The mGluR2/3 inhibition is always faster than the mGluR1 excitation and therefore suppress firing transiently prior to excitation. The mGluR2/3 currents also systematically vary in time course, but interestingly, and in contrast to the mGluR1 excitation, these currents are longer lasting at the extreme in cells lacking mGluR1, and they become faster in cells where mGluR1 is present. Consequently, MF stimulation evoked pauses are briefer in UBCs that have subsequent shorter-lived increases in firing.

### General implications for other cell types

It is likely that similar mechanisms are at play for generating continuous gradients in the amplitude and time course of Group I mGluR-dependent signaling in other brain regions, because we found similar correlated gradients of mGluR1/5, PLCβ, and DGKβ/γ in layer 2/3 cortical neurons and in Purkinje cells (**Figure 1a, c**). Although we have focused on the excitatory TRPC3 mediated currents in UBCs, mGluR1/5 signaling extends to other aspects of synaptic function, including retrograde signaling by endocannabinoids, and regulation of long-term synaptic plasticity^34,35^. This raises the intriguing possibility that these processes are regulated in a continuous manner in cell types with correlated gradients of expression of mGluR1/5 signaling.

### Cell-intrinsic neural integrators

Our results suggest that transcriptomic control of metabotropic signaling generates a family of smoothly varying temporal transformations. The correlated change across numerous response properties suggests a remarkable level of coordination between multiple post-synaptic molecular drivers. This cell-intrinsic synaptic mechanism for generating long-lasting responses is not prone to noise accumulation or catastrophic failure, as can occur with network-based mechanisms^36-38^. The transient firing suppression is reminiscent of the stimulus-onset quenching of neural variability observed in many cortical neurons^39^, and could mediate a ‘soft-reset’ function at stimulus onset, which is particularly useful for controlling noise accumulation in cells with long integration time constants^38^.

### Optimal basis set for temporal learning

Certain features of the basis expansion by UBCs likely have important implications for cerebellar learning. On a logarithmic time axis, UBC responses were well-approximated by shifted Gaussians (**Figure 6b,j**, schematized in **Figure 7d**). By direct analogy to radial basis functions, the UBC basis set may allow the cerebellum to model arbitrary input-output functions between hundreds of millisecond to tens of seconds^40,41^. Crucially, the fact that early responding UBCs had narrow temporal tuning curves while late responding UBCs had wide tuning curves (**Figure 6i, j**) suggests that the precision of output generated from this basis set should scale inversely with the time delay. This may provide a cellular substrate for Weber’s Law, a linear correlation between mean and variability of response, frequently observed in cerebellar dependent timing tasks^42,43^. Representation and learning below the hundred-millisecond range is likely supported conjunctively by several synaptic mechanisms^44^. Lastly, the UBC basis set provides a dense temporal representation as individual responses overlap significantly in time. This is in contrast to sparse temporal representations such as the sequential burst of HVC neurons in bird songs^45^. Unlike in the imitation learning of songs, the cerebellum undergoes life-long adaptations. Therefore, the utilization of dense versus sparse temporal representations may reflect the underlying tasks, and the degree of generalization versus memorization required by the behavioral task. By demonstrating that a molecularly diverse type of neuron generates a multi-scaled basis expansion that is robust and flexible, our results highlight the functional relevance of molecular heterogeneity in central circuits.

## Methods

### Ethics

All animal procedures were carried out in accordance with the NIH and Animal Care and Use Committee (IACUC) guidelines and protocols approved by the Harvard Medical Area Standing Committee on Animals (animal protocol #1493).

### scRNA-seq

scRNA-seq data for UBC and Purkinje cells was obtained in Kozareva et al.2020^6^. In brief, high-throughput single-nucleus RNA-seq (snRNA-seq) was used to analyze 780,553 nuclei isolated from 16 different lobules from 6 p60 mice. Of these, 1613 were categorized as UBCs and 16634 as Purkinje cells. Although UBCs were found in all lobules, their density was regionally dependent, and as is well-established the density was particularly high in regions involved in vestibular processing such as lobules IX and X. scRNAseq data for L2/3 cortical neurons was obtained from Yao et al., 2020^46^.

### Slice preparation

Cerebellar slices were prepared from adult (P30-40) C57BL/6 mice of either sex. Animals were anesthetized by a peritoneal injection of 100/10 mg/kg ketamine/xylazine mixture and then intracardially perfused with ice-cold cutting solution (in mM): 110 Choline Cl, 2.5 KCl, 1.2 NaH_2_PO_4_, 25 NaHCO_3_, 25 glucose, 0.5 CaCl_2_, 7 MgSO_4_ 2, 2.4 Na-pyruvate, 11.6 Na-ascorbate and 25 glucose equilibrated with 95% O_2_ and 5% CO_2_. Cerebellum was subsequently extracted, submerged in the cutting solution and 250μm sagittal slices from the vermis were obtain using a Leica VT1200S vibratome (Leica Biosystems Inc. Buffalo Grove, IL). Slices were transferred into an incubation chamber with artificial cerebral spinal fluid (ACSF) containing (in mM): 125 NaCl, 26 NaHCO_3_, 1.25 NaH_2_PO_4_, 2.5 KCl, 1 MgCl_2_, 1.5 CaCl_2_ and 25 glucose and equilibrated with 95% O_2_ and 5% CO_2_ (pH 7.4, osmolarity 315). Following 30 minutes of incubation at 32°C, the slices are kept for up to 5 hours at room temperature.

### Electrophysiology

Recordings were performed under physiological temperature (34~36°C) in ACSF containing inhibitory receptor blockers 20μM picrotoxin and 1μM CGP. Optically guided cell-attached or whole-cell recordings were made in lobule X under an Olympus BX51WI microscope equipped with differential interference contrast (DIC) components. Patch-pipette of (2~3 MΩ for cell-attached recording, 4~5 MΩ for whole-cell recording) were pulled from borosilicate capillary glass (World Precision Instruments) with a Stutter P-97 horizontal puller. Mossy fiber (MF) stimulations were delivered via a bipolar theta glass pipette filled with bath solution positioned at least 50 μm away to avoid stimulating the recorded neuron. Care was taken during the search of the sole MF input to elicit reliable response and avoid antidromic spikes. Cell-attached recordings were made using ACSF with a loose patch of 100 MΩ. It is important to avoid using high-potassium internal solution for cell-attached recording as we have noticed significant depolarization block in persistently firing cells. In the case where both cell-attached and whole-cell recordings were done on the same cell, extracellular electrodes were swapped out with a second electrode with an internal solution containing (in mM): 122 K-methanesulfonate, 9 NaCl, 9 HEPES, 0.036 CaCl_2_, 1.62 MgCl_2_, and 0.18 EGTA. A separately aliquoted regenerating solution containing: 4 Mg-ATP, 0.3 Tris-GTP, 14 Tris-creatine phosphate was kept in a −20°C freezer and added to the internal alone with 10 μM Alexa 589 dye on the day of recording (pH 7.4 and osmolarity 315). A - 8mV liquid junction potential was corrected between the internal and external solutions. Series compensation was not made for voltage-clamp recording. Data were collected with Multiclamp 700B amplifier (Molecular Devices, LLC., San Jose, CA), filtered at 4kHz (4-pole Bessel filter) online, digitized at 50kHz with ITC18 (Heka Instrument, Inc., Holliston, MA), and saved using custom software (courtesy of Matthew Xu-Friedman, SUNY Buffalo, Buffalo, NY) in Igor Pro (WaveMetrics Inc., Portland, OR) for offline analysis.

### Pharmacology

Sequential drug wash-ins were performed using a computer-controlled solenoid manifold system (ValveLink 8.2, Automate Scientific, Inc., Berkeley, CA) with a flow rate of 1 to 2mL/min, where indicated, with mGluR1 antagonist (100μM LY357385, Tocris Bio-Techne, Minneapolis, MN), AMPA receptor blocker (5μM NBQX), NMDA receptor blocker (2 μM R-CPP), mGluR2/3 antagonist (1μM LY341495, Tocris Bio-Techne, Minneapolis, MN) and DGK inhibitor II (100 μM R59949, MilliporeSigma, St. Louis, MO)

### One-photon glutamate uncaging

Slices were incubated in 200 μM RuBi-Glutamate ACSF solution in the presence of inhibitory blockers 20μM picrotoxin and 1μM CGP at physiological temperature (34~36°C). The osmolarity of the solution was monitored and adjusted with deionized water. After measuring the response to MF stimulation, optical uncaging of glutamate was done by 20×100Hz pulse train with 2ms to 4ms pulse width illumination over a 20μm diameter spot above the dendritic brush. The light stimuli were delivered by a blue laser (MBL-III-473-50mW, Optoengine, Midvale, UT) through a 60x objective at >160mW/mm^2^.

### Data analysis

Pharmacology experiments were reported as time series plots showing mean±sem of spiking or current response during drug wash-in. Spiking response properties such as pause time and baseline firing were analyzed on average instantaneous firing rates over three to five trials. Measurements of half-decay time, amplitude, and width are extracted from a smooth response based on log-Gaussian fitted to the instantaneous firing rate using the nonlinear curve fitting toolbox in MATLAB. The parameterization used for log-Gaussian fit is 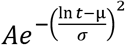, where *A* is amplitude, *μ* is the log peak location and *σ* is the width of temporal response on a logarithmic time scale. All linear and non-linear regressions are performed in MATLAB and reported with adjusted *R*^2^ values, calculated as 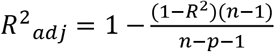, where *n* is the sample size, p is the number of model parameters, and *R*^2^ is calculated as 1 minus the ratio of residual sum of squares over the total sum of squares, to assess the quality of the fits. An inverted soft-plus function of the form 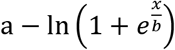 was used to fit the log-transformed amplitude values vs. cell ID in **Figure 6h**. Kolmogorov-Smirnov test was used for testing whether the distribution of peak time followed a uniform distribution on a logarithmic timescale.

The population response amplitude was defined as the L_2_-norm of the population activity vector, which each dimension being the instantaneous firing rate of one UBC over time. For ease of interpretation and to control for the expected 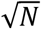 increase in population amplitude as a function of N, the number of measured neurons, the population amplitude was normalized by 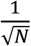 in **Figure 6**. Population response were fitted with a shifted power law of the form *Cn*(*t + t_min_n^β^*)^∝^, where *n* is the number of stimuli, *C* is a scaling constant for amplitude, *t_min_* and *β* control when and how fast power-law behavior dominates for different numbers of stimuli and *∝* is the critical exponent.

## Data Availability

The datasets are available from the corresponding author on reasonable request.

## Code Availability

The custom code used in the analysis will be available from Chong Guo’s GitHub repository upon publication: https://github.com/chongguo.

