## Extended Data Figures for "Graded heterogeneity of metabotropic signaling underlies a continuum of cell-intrinsic temporal responses"

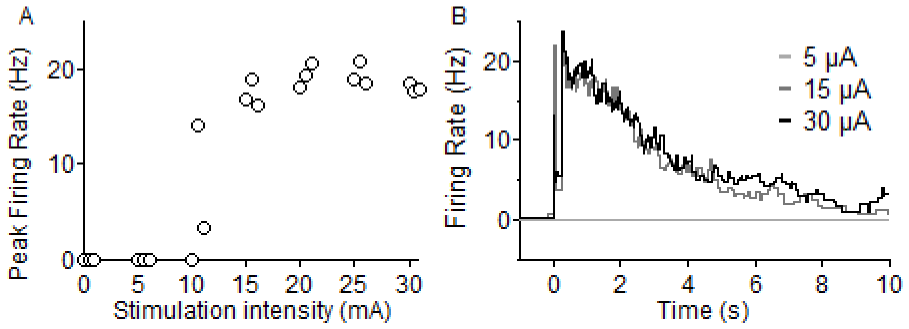


**Extended Data Figure 1. Varying stimuli intensities did not alter temporal profiles of spiking responses in UBCs**

A. Electrical stimulation (20x100Hz) evoked all-or-none response as shown in the peak firing rate of sample UBC, three trials are done at each intensity. Unreliable responses are observed at an in intermediate intensity (10 µA) likely due to failed axonal stimulation.
B. The decay kinetics of the evoked response did not depend on the stimulation intensity past the threshold required for reliable response (15 µA).

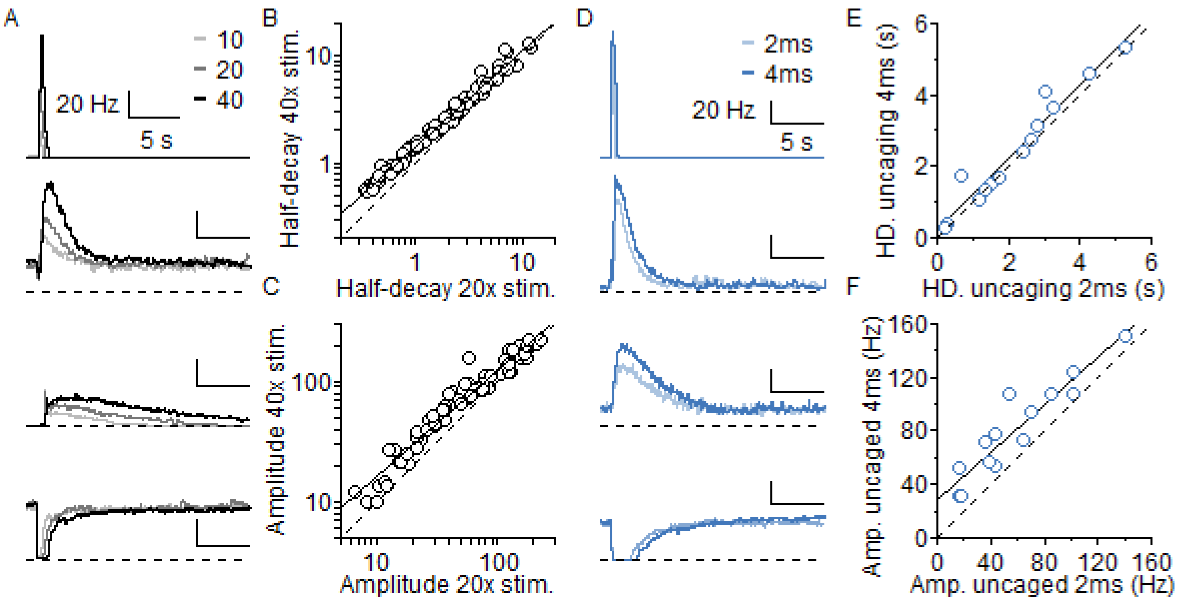


**Extended Data Figure 2. Diversity of spiking responses in UBCs is not a result of variations in stimulation parameters.**

A. Examples of instantaneous firing rate curves for four sample UBCs to a 100 Hz burst of MF inputs consisting of 10 (light grey), 20 (grey), and 40 (dark) stimuli. The dashed lines are at 0 Hz.
B. Half-decay time of spiking response for 20 vs. 40 stimuli across all cells (black marker), linear fit on the $\log_{10}$ transformed variables (solid line, $R_{adj}^2$=0.98, slope=0.89, intercept=0.15) and the unit line (dotted line).
C. Peak amplitude of spiking response for 20 vs. 40 stimuli across all cells (black marker), linear fit on the $\log_{10}$ transformed variables (solid line, $R_{adj}^2$=0.95, slope=0.88, intercept=0.336) and the unit line (dotted line).
D. Examples of instantaneous firing rate curves in the glutamate uncaging experiment for four sample UBCs to a 100 Hz burst of 20 light flashes with pulse durations of either 2ms (light blue) or 4ms (dark blue). The dashed lines are at 0 Hz.
E. Half-decay time of spiking response for 2ms vs. 4ms pulse across all cells (blue marker), linear fit (solid line, $R_{adj}^2$=0.93, slope=2.03, intercept=0.07) and the unit line (dotted line).
F. Peak amplitude of spiking response for 2ms vs. 4ms pulse across all cells (blue marker), linear fit (solid line, $R_{adj}^2$=0.86, slope=0.89, intercept=29.02) and the unit line (dotted line).

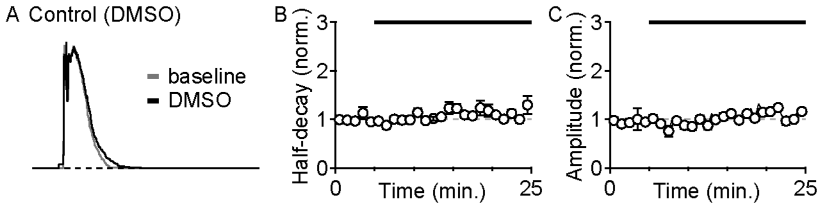


**Extended Data Figure 3. Summary of MF-evoked spiking response for control (DMSO) drug application under cell-attached configuration.**

A. Example of instantaneous firing rate before (gray) and after DMSO wash-in (black).
B. Summary of half-decay time of instantaneous firing rate response with DMSO (normalized to baseline, mean±sem, n=6).
C. Summary of peak amplitude of instantaneous firing rate response with DMSO (normalized to baseline, mean±sem, n=6)

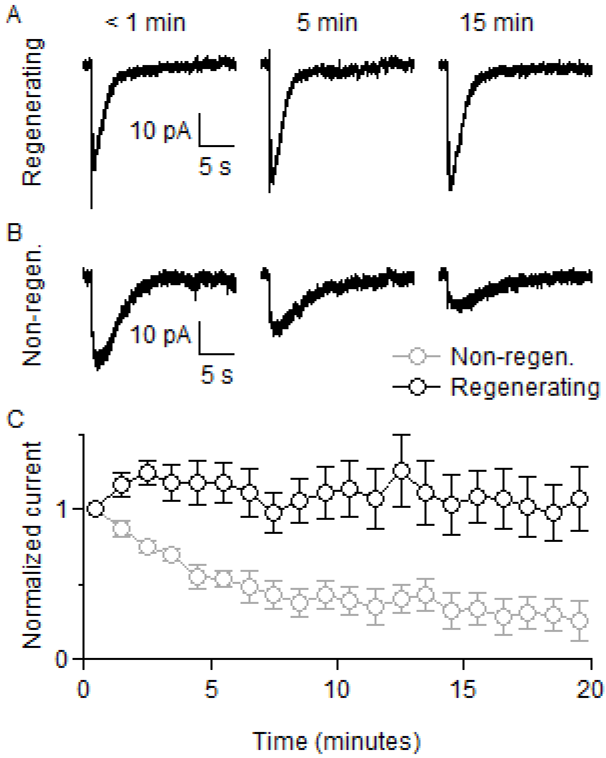


**Extended Data Figure 4. Synaptic currents in UBC washed out over time without a regenerating internal solution**

A. Synaptically-evoked currents (20x100 Hz) are shown less than 1 min (left), 5 min (middle), and 15 minutes (left) after break-in, for a non-regenerating internal solution.
B. Same as in A but for recordings using a regenerating internal solution.
C. Summary of responses over time (normalized to fist point, mean ± sem, non-regenerating internal in grey n = 5, regenerating internal in black n=6)

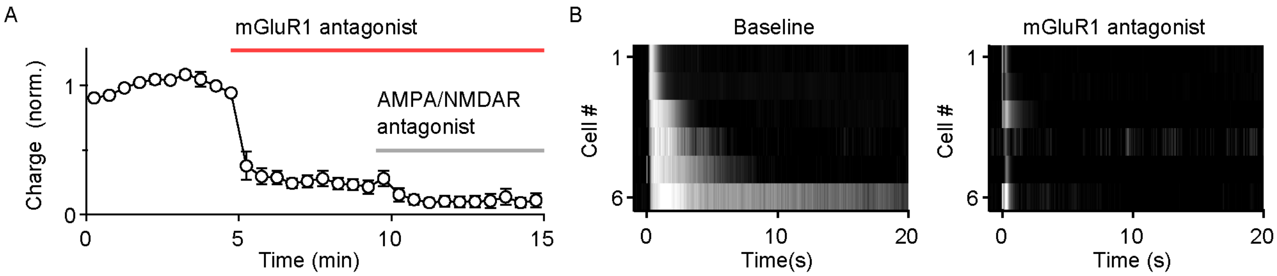


**Extended Data Figure 5. Sequential applications of mGluR1 and AMPA/NMDA receptor antagonists revealed diverse kinetics of mGluR1-mediated synaptic currents.**

A. Summary of evoked synaptic charge (20x100 Hz) and the effect of an mGluR1 antagonist (red bar), followed by the co-application of antagonists of mGluR1, AMPA and NMDA receptors (grey bar). Amplitudes are normalized to baseline (mean±sem, n=6).
B. Heatmap of the whole-cell recordings of current responses before (top) and after (bottom) mGluR1 antagonist wash-in. Responses are normalized to the peak current responses measured before drug application.

**Extended Data Figure 6. Sample UBC spiking responses to burst and rate modulated MF input.**

Most of our experiments focused on the response evoked by bursts of MF stimuli. However, UBCs lobule X are often studied in response to sinusoidal MF stimulation (Zampini et al., 2016). We therefore characterized burst and sinusoidal modulations of MF input in the same cells.

A. Representative instantaneous firing rate of a fast (left) and a slow (right) UBC with burst MF stimulations (20x100 Hz). Each trace is an average of 4 trials

B. Sine-wave modulated firing rate of MF around 30 Hz baseline with 30 Hz amplitude. Each trace is an average of 4~50 trials

C-G. Instantaneous firing rate of the same fast (left column) and slow (right column) UBCs to 0.2, 0.5, 1, 2 and 5 Hz sine-wave modulated MF input

H. Phase-delays (measured in 0.5 Hz condition) under sine-wave modulated MF stimulations correlate with the half-decay times of burst stimulation responses (solid line, $R_{adj}^2$=0.57, slope=0.58, intercept=0.71, n = 9).

We found a positive correlation between the half-decay time of the burst response and the phase delay of the sinusoidal response. This suggests that under physiologically relevant conditions, diverse mGluR1-dependent temporal kinetics of burst response is related to diverse phase response in the frequency domain.

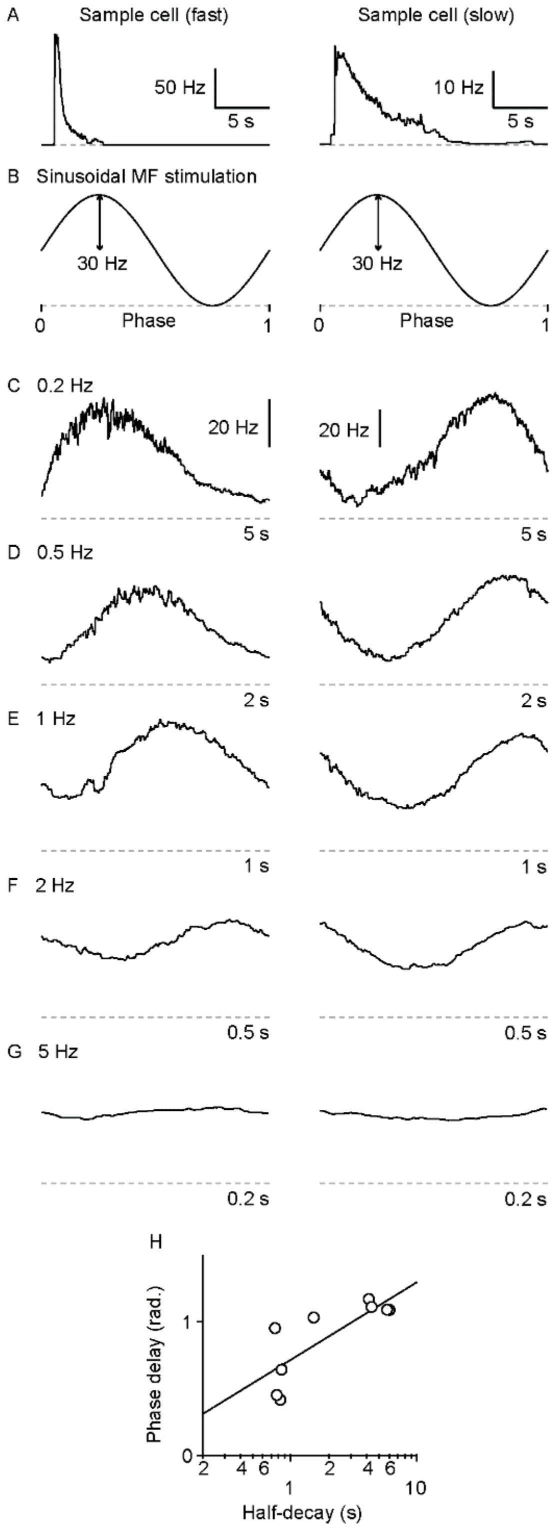
